# ‘Synergistic-cidal’ effect of amoxicillin conjugated silver nanoparticles against *Escherichia coli*

**DOI:** 10.1101/832568

**Authors:** Arivarasan Vishnu Kirthi, Loganathan Karthik, Jayachandran Venkatesan, Atul Changdev Chaskar

## Abstract

We are reporting the synergistic effects of amoxicillin coated silver nanoparticles, and sodium cholate is used as the stabilizer. Following, the synthesis of AgNPs physicochemical characterization was carried out using UV spectroscopy, XRD, FTIR, FESEM, AFM and NC-AFM analysis. The confirmation of AgNPs was confirmed UV spectral peak at 426 nm, the morphology of the nanoparticles was found to be spherical, oval with an average size of 35.50 nm. The cytotoxic effect was carried out using MTT assay with IC50 of 50.78 µg/mL and the antibacterial effect was observed as zone of inhibition of 12 mm for E.coli. The drug release was determined as 69%, which contributes to the therapeutic efficacy of water soluble based nanoparticles/nanomedicines for biomedical applications.

## 1. Introduction

Recent advances in biotechnology have accelerated the discovery of novel drug molecules. Metal-based nanoparticles possess several advantages over other systems, such as cell specific targeting of the nanoparticle delivery system and the ability to control the kinetics of drug release [1]. Silver (I) oxide takes advantage of their low solubility in aqueous environments to slowly release silver ions as Ag (OH)^−2^. It is well known that metallic silver is complexes by silver ions rendering silver clusters (Ag^2+^,Ag_4_^2+^) [2]. Consequently, there is a need to produce virtuous, non-toxic and environment friendly methods for the synthesis of AgNPs. The utilization of the biological system in this area is rapidly gaining importance due to its growing success.

Amoxicillin trihydrate (α-amino-hydroxybenzyl-Penicillin) is a semi-synthetic, orally absorbed and broad-spectrum antibiotic. The monodispersed AgNPs with narrow particle size distribution were most effective antibacterial agents because of their high surface/volume fraction so that a large proportion of silver atoms were in direct contact with their environment [3].

The facile one pot synthesis of amoxicillin and sodium salt of amoxicillin stabilized gold nanoparticles has been reported [4]. Chitosan/polygamma glutamic acid nanoparticles incorporated into pH-sensitive hydrogels were developed as an efficient carrier for amoxicillin delivery specifically with intercellular spaces against *Helicobacter pylori* infection [5]. Polyethylene glycol coated polyethylcyanoacrylate nanoparticles loaded with amoxicillin were prepared and the influence of the PEG coating on the particle size, zeta potential, drug release rate and phagocytic uptake by murine macrophages [6]. Due to the emergence of antibiotic resistant bacteria and limitations of the use of antibiotics, the clinicians have returned to silver wound dressings containing varying levels of silver [7].

The synthesized AgNPs prepared on polydimethylsiloxane (PDMS) for the spatial control of cell capture, where the residual Si-H groups in the PDMS matrix were used as reductants to reduce AgNO_3_ for the formation of AgNPs and PDMS-AgNPs composite showed good antibacterial property against *Escherichia coli* [8]. A number of different species of bacteria and fungi are able to reduce metal ions producing metallic nanoparticles with antimicrobial properties [9, 10]. Recently, efficient antibacterial activity was observed against multidrug resistant and highly pathogenic bacteria, including multidrug resistant *Staphylococcus aureus*, *S*. *epidermidis*, *Salmonella typhi* and *E*. *coli* by AgNPs produced by the fungus *Fusarium acuminatum* [11].

In the present work, we report synthesis of nanoparticles that allows us to stabilize AgNPs with capping agents featuring amoxicillin and sodium cholate, in order to investigate their use as antibacterial agents. NPs are prepared with a well-established method that yielded reproducible amoxicillin capped AgNPs of defined dimensions and shapes. For averting particle aggregation and imparting stability of AgNPs, the surfactant sodium cholate was used. Amoxicillin is an organic molecule having two potential functional groups, amide and carbonyl and may be suitable for synthesis of AgNPs as a stabilizer or surfactant. Finally, it is described how these biomimetically synthesized nanoparticles can be used for investigation of their antimicrobial activity against bacteria.

## 2. Material and Methods

### 2.1. Experimental section

The silver amoxicillin complex nanoparticles were synthesized by the slightly modified methods [12, 13]. Briefly, 10 mL of 1M amoxicillin solution and 50µl of sodium cholate (pH 11; adjusted by adding sodium hydroxide) was mixed with 30 mL of 1 M aqueous AgNO_3_ solution. The formation of silver amoxicillin nanoparticles with pH 6 was accompanied by a very rapid formation of white precipitate. The dispersion was then centrifuged for half an hour at 12, 000 rpm at room temperature to form surface modified AgNPs. The pellet was washed three times with double distilled water, and finally dispersed in 50 mL of phosphate-buffered saline (PBS) buffer (pH 6.0) by sonication in cold condition. It was further purified by centrifugation at 4°C for 2 h to remove unbound sodium cholate.

### 2.2. Characterization of AgNPs

The bioreduction of Ag^+^ ion in solution was scrutinized by UV-visible spectrometer (Techomp 8500 spectrometer). Further, the characterization was completed using FTIR (Bruker tensor 27) spectrometer. In order to remove any free biomass residue, the residual solution was centrifuged at 5000 rpm for 30 min and the resulting suspension was redispersed in 10 mL sterile distilled water. The dried samples were pelletized with KBr and analyzed using FTIR. X-ray diffraction was performed using Bruker, DK advance, Germany operated at 30 kV and 100 mA. The pattern was recorded by CuKα radiation with λ of 1.5406 Å and nickel monochromator. The scanning was done in the region of 2θ from 10° to 90°. The size of the nanoparticles was calculated through the Scherer’s equation [14].

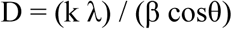

Where D is the average crystal size, k is the Scherer’s coefficient (0.89), λ is the X-ray wave length (λ=1.5406 Å), θ is Bragg’s angle (2θ), and β the full width at half maximum (FWHM) in radians.

The synthesized AgNPs were prepared at the weight ratio of 2:1 (drug/polymer). For electron microscopic studies, 25 μL of sample was sputter-coated on copper stub, and the images of nanoparticles were studied using Field-emission scanning electron microscopy (FESEM, JSM-6700, JEOL, Japan). Topography was studied using Atomic force microscope (AFM and NC-AFM mode) (Veeco Pico Force) working in the contact mode. A thin film of the sample was prepared on a glass slide by dropping 100 µL of the sample on the slide, and was allowed to dry for 5 min. Characterization was done by observing the patterns on the surface topography and data analysis through WSXM software [15].

### 2.3. Agar well-diffusion method

The bacterial pathogen *E*. *coli* (multi drug resistant strain) was obtained from the MTCC, Chandigarh, India (MTCC-443). The strain was maintained on Nutrient agar slant at 4°C.MHB plates were swabbed on three axes with a sterile cotton-dipped swab in the freshly prepared diluted culture. The holes were filled with 100 µl of different concentrations of AgNPs (50, 100, 150, 200 and 250 µg/mL) and allowed to stand for 1 h. The plates were kept for further incubation at 30°C for 24 h. Deionized water was used as a negative control and the antibiotics amoxicillin (30 µg/0.1 mL) used as positive control [16].

#### 2.3.1. Minimum inhibitory concentration (MIC)

MIC was determined according to the standard broth micro-dilution method [17]. The initial concentration of the nanoparticles showed minimum antimicrobial activity in the agar well-diffusion method was serially diluted to obtain different concentrations. The culture broths of the test organisms were diluted to contain ~10^5^ to 10^6^ CFU/mL.

### 2.4. In vitro antioxidant assay

#### 2.4.1. DPPH radical scavenging activity

The DPPH free radical scavenging assay was conducted as per the method of [18]. The different concentrations of AgNPs were prepared (10, 20, 40, 60, 80 and 100μg/mL) with double distilled water. 2 mL of each dilution was mixed with 1 mL of DPPH solution (0.2 mM/mL in methanol) and mixed thoroughly. The mixture was incubated in dark at 20°C for 40 min. Absorbance was measured at 517 nm using UV-visible spectrophotometer with methanol as blank. Gallic acid was used as positive control. The percentage of scavenging activity of DPPH of the AgNPs was calculated according to the following formula:

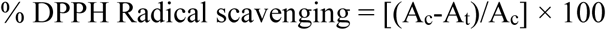

Here, A_c_ is the absorbance of the control (DPPH), A_t_ is the absorbance of test sample [19].

#### 2.4.2. Reducing power activity

The reducing power of AgNPs was determined by ferric reducing power assay [20]. 1 mL of synthesized AgNPs at different concentrations (125, 250, 500 and 1000 μg/mL) were mixed with phosphate buffer (2.5 mL, 0.2 M, pH 6.6) and 2.5 mL of 1 % potassium ferricyanide (K_3_Fe(CN)_6_. The mixture was incubated at 50°C for 20 min. A volume of 2.5 mL of trichloroacetic acid (10%) was added to the mixture, and centrifuged at 3000 rpm for 10 min in a cooling centrifuge. 2.5 mL of supernatant was mixed with equal volume of distilled water and 0.5 mL FeCl_3_ (0.1%). Absorbance was measured at 700 nm using UV-visible spectrophotometer. Ascorbic acid was used as positive control. Higher absorbance of the reaction mixture indicated greater reductive potential. Each experiment was performed in triplicates at each concentration

#### 2.4.3. Total antioxidant activity

The total antioxidant activity was measured by modified method of Prieto, Pineda [21]. The reaction mixture of 3 mL with 0.6 M sulphuric acid, 28 mM sodium phosphate and 1% ammonium molybdate and added to the different concentrations of AgNPs (10, 50, 100, 250 and 500 mg) and kept in a water bath at 95°C for 60 min. Absorbance was recorded at 695 nm, ascorbic acid was used as the standard and the total antioxidant activity was expressed as equivalents of ascorbic acid.

### 2.5. Anti-cancer activity of AgNPs against Hep-G2 cancer cell lines

#### 2.5.1. Synthesized AgNPs preparation

The AgNPs was dissolved in different concentrations ranging from 25 to 500µg/mL in 10 % DMSO and did not affect cell survival. AgNPs was dissolved in different concentrations ranging from 25, 50, 100, 200 and 500µg/mL in 1.0 % DMSO. The 25 µg/mL was found to cause no visible changes in the cellular morphology of the cell line, the plaque formation and the cell necrosis occurred from 50µg/mL.

#### 2.5.2. MTT assay

The viability of cells was assessed by MTT assay using Hep-G2 cell lines. 3-(4,5-dimethylthiazol-2-yl)-2,5-diphenyl tetrazolium bromide was dissolved in Dulbecco’s PBS (-) (Sigma-Aldrich, India) (pH 7.4) at 5 mg/mL and filtered to sterilize and remove a small amount of insoluble residual used in MTT assay [22]. The Hep-G2 cells were plated separately in 96 well plates at concentration of 1×10^5^ cells/well. After 24 h, cells were washed twice with 100 µl of serum-free medium and starved for an hour at 37°C. After starvation, cells were treated with different concentrations of test AgNPs (25-500 µg/mL) for 24 h. At the end of the treatment period the medium was aspirated and serum free medium containing MTT (0.5 mg/mL) was added and incubated for 4 h at 37 ºC in CO_2_ incubator. 50% inhibitory concentration value (IC_50_) of the synthesized AgNPs was identified for normal fibroblast cell line [22, 23].

The MTT containing medium was discarded and the cells were washed with PBS (200 µl). The crystals were then dissolved by adding 100 µl of DMSO and mixed properly by pipetting up and down. Spectrophotometric absorbance of the purple blue formazan dye was measured in a microplate reader at 570 nm (Biorad 680). Cytotoxicity was determined using Graph pad prism 6 software [24]. The viability was calculated using the following formula:

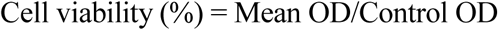

#### 2.5.4. In vitro drug release kinetics

5 mg of AgNPs were suspended in 2 mL phosphate buffered saline (PBS, 0.1 M, pH 7.4) and sonicated for 30 sec at 15 watts by using Ultrasonic homogenizer, UH-50, SMT Co. Ltd., Japan and transferred into a dialysis bag (Mw =12 KDa). The dialysis bag was placed in to 100 mL bottle with 50mL PBS and the medium was stirred at 100 rpm at 37°C. The media was changed to study the drug release kinetics and prevented the saturation of the drug. The concentration of the released AgNPs into PBS was determined by UV-visible spectrophotometer (Schimadzu UV-1601) at 310 nm. A continuous release of AgNPs was measured over a period of 10 h. The percentage of silver released was calculated using the equation:

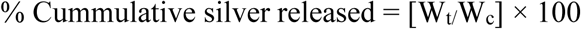

Where, Wc is the total silver content in the dialysis bag and Wt is the silver content in the PBS medium at a time [25, 26]. The synthesized AgNPs with amoxicillin were tested against *E*. *coli*.

## 3. Results and discussion

### 3.1. Possible mechanism for the synthesis of the AgNPs

The formation of complex between AgNPs and amoxicillin was confirmed by using Fourier transform infrared spectroscopy spectra. The spectrum of AgNO_3_ showed the sharp peak at 1379 cm^−1^ was from the presence of NO_2_ group. The spectrum of amoxicillin showed broad peak at 3462 cm^−1^ was assigned to the presence of hydroxyl group. Prominent and very sharp peaks at 1775 and 1686 cm^−1^ was concluded due to presence of carbonyl (C=O) groups. The spectrum for the synthesized AgNPs clearly showed the formation of complex between amoxicillin and AgNPs, due to the absence of C=O peaks at 1775 and 1686 cm^−1^ and OH group at 3462 cm^−1^. The present results revealed that the FTIR spectra showed a complex formation between AgNPs and amoxicillin drug. FTIR of AgNPs conjugated amoxicillin, amoxicillin, and AgNO_3_ 1mM were shown in Fig. S1.

#### 3.1.1. UV spectrophotometry study

UV-Vis absorption spectra demonstrated a novel procedure for the preparation of the monodispersed nanoparticles was shown in Fig. 1A. An absorption peak was found at 426 nm for the AgNPs and the control was at 420 nm (3.32 eV).

**Fig 1.**
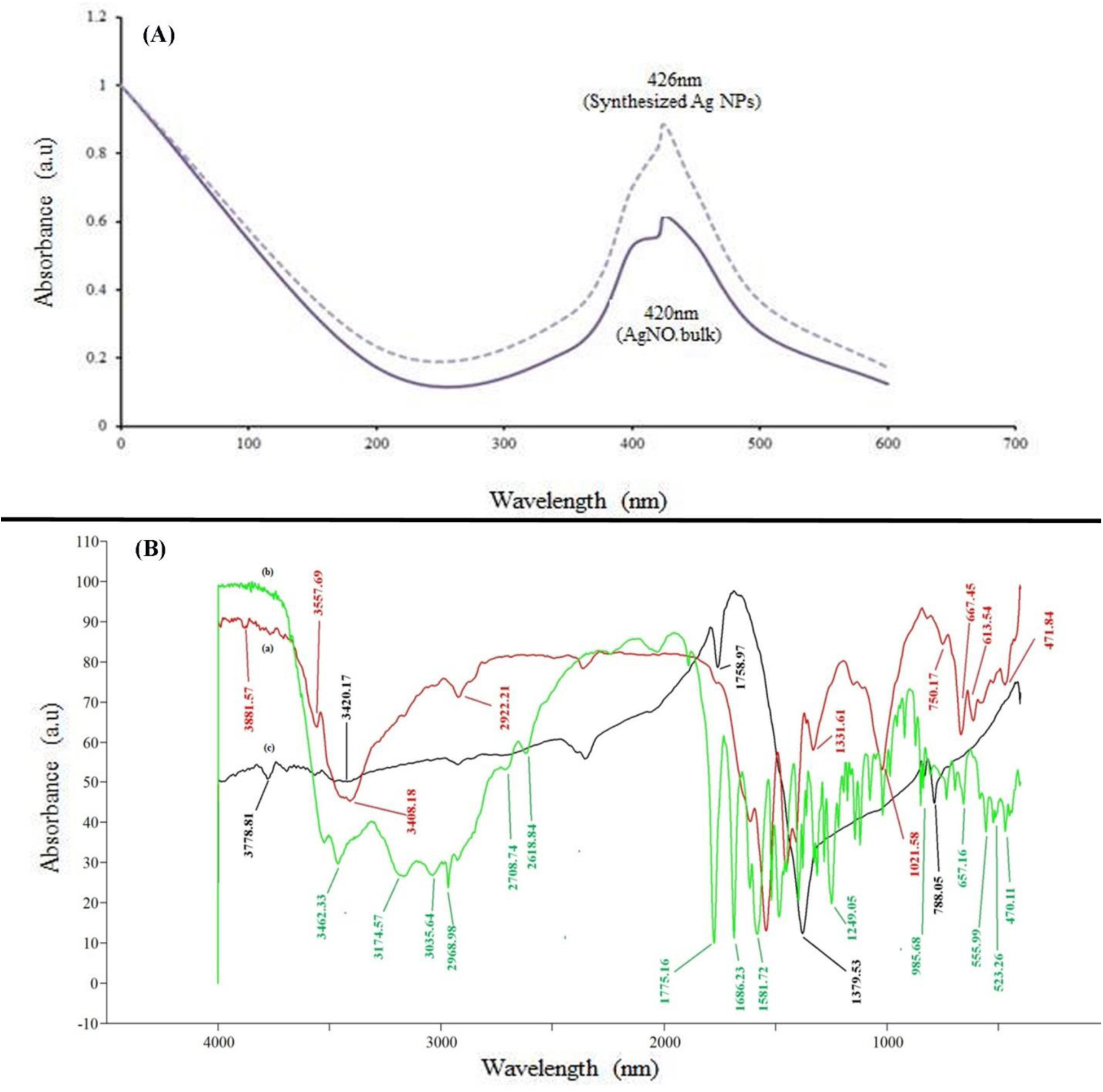
(A). UV absorption spectrum for the synthesized Ag NPs functionalized amoxicillin showing peak at 426 nm and control silver nitrate (bulk) at 420nm. (B). FTIR for the Ag NPs with amoxicillin (red line-a), amoxicillin drug as control (green line-b) and silver nitrate 1mM control (black line-c).

#### 3.1.2. Fourier transform infrared spectroscopy (FTIR)

The Fourier transform infrared spectroscopy for the AgNPs (red line), amoxicillin control (green line), and AgNO_3_ 1mM control (black line) are shown in Fig. 1B. The final product contains the functional groups of both the silver and amoxicillin, showed the confirmation of the synthesis AgNPs. These functional groups included intermolecular bonded (strong), presence of mononuclear benzene aromatics, ketone groups, carboxylic groups, intermolecular H-H bonds, aldehyde functional structures, aryl-C-CH_2_, σ-hydroxyl aryl ketone, ketone, anhydrides and N-H oscillations, which were corresponding to the frequency numbers at 3462.23, 3174.55, 3035.64, 2968.98, 2708.74, 2618.84, 1775.16, 1686.23, 1581.72, and 555.99 cm^−1^, respectively. The spectrum amoxicillin (control) was found to be the homologous functional groups in the drug. The functional groups were intramolecular bonds (weak and strong) (3881.57 cm^−1^), intermolecular bonds of H-H (3557.89 cm^−1^), alkynes (3408.18 cm^−1^), alkenes (1331.61 cm^−1^), mononuclear aromatics (1155.96 cm^−1^), amides (750.17 cm^−1^) and primary amides (667.14 cm^−1^). In the case of AgNPs, there were no changes in the absorption frequencies of other groups except the 1775.16, 1686.23 and 1581.72 cm^−1^ which are corresponding to various ketone subgroups-acyclic, and aryl ketone. From the above data, it may be concluded that neither the NH_2_, nor group bound to the silver surface. It is only the nitrogen atoms of the free amino group that binds directly on the silver exterior, and the carbon atoms of the free ketone group that binds directly on the silver surface.

FTIR analysis characterized the formation of complex between amoxicillin and AgNPs. The absence of nitrate group in the product spectrum showed the formation of AgNPs. The spectrum for the amoxicillin functionalized AgNPs clearly shows the formation of complex between amoxicillin and AgNPs, due to the absence of C=O peaks at 1775 and 1686 cm^−1^ and OH group at 3462 cm^−1^ and it was found similar to previous reports [27, 28].

#### 3.1.3. X-ray diffraction (XRD) analysis

X-ray diffraction analysis showed the peaks at 17.509° (002), 28.844° (121), 29.617° (211), and 38.439° (123) which is in corresponding to the JCPDS file no. 84-0713. A mathematical analysis of the Bragg’s peaks was undertaken to calculate the crystallite size using the Scherrer’s formula, D = kλ / β cos θ where D is the crystalline size, k is a constant (the particles were spherical in shape, 0.89), λ is the wavelength of the X-ray radiation, β is the line width (obtained after correction for the instrumental broadening) and ‘θ’ is the angle of diffraction for AgNPs (Figs. 2 A and B). The average particle size obtained from XRD data was found to be about 35.50 nm. We ascribed the discrepancy to the polydispersity of our samples, a well-known interference factor in classical XRD data interpretation of nanomaterials [29]. Additionally, the peaks attributed to silver-*fcc* in present case showed peculiar shoulders, suggesting that the structure of the metallic colloid is very complex at the nanoscale [30].

**Fig 2.**
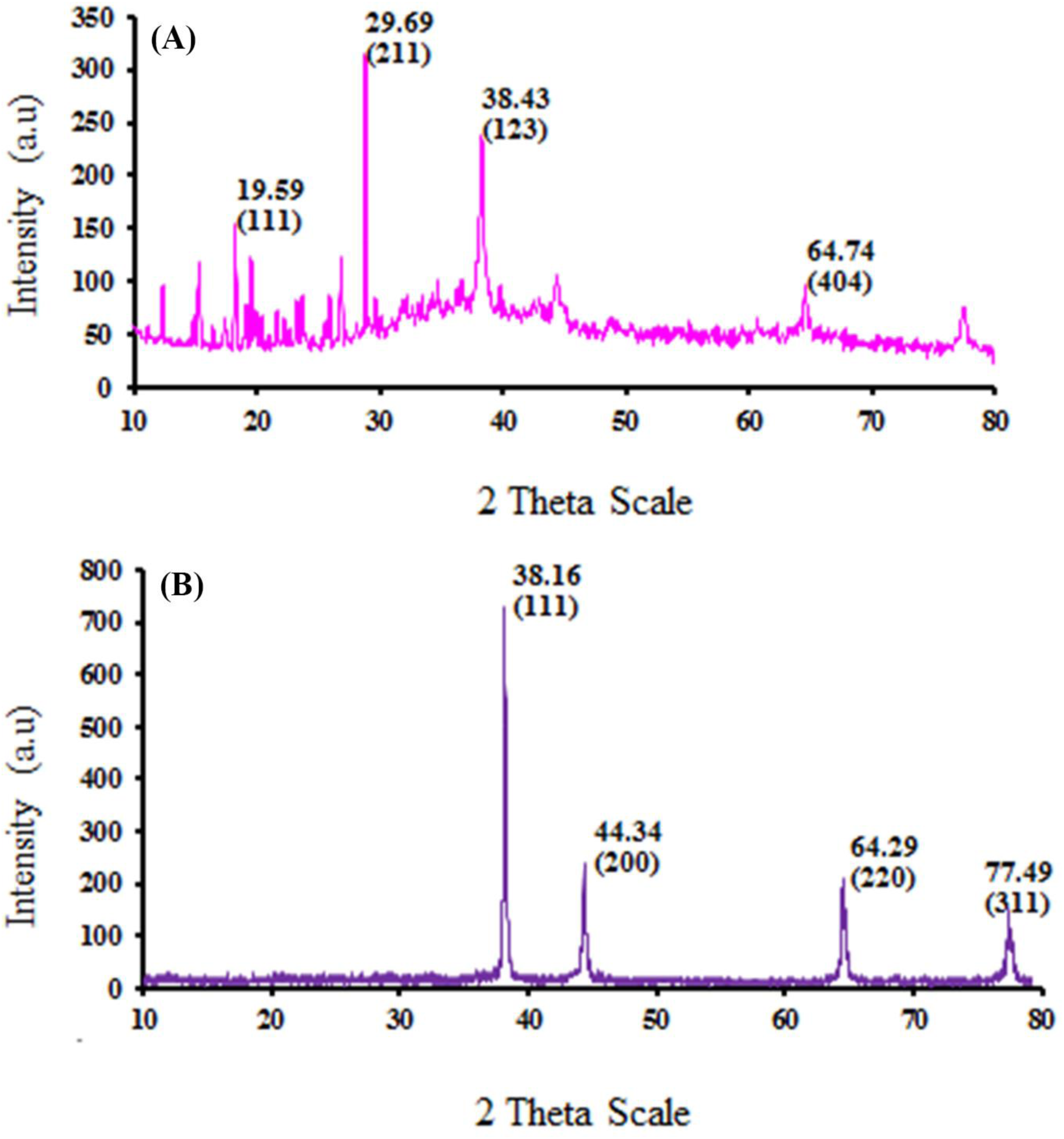
Indexed XRD of (A) Ag NPs with amoxicillin at room temperature (B). Silver (bulk) spectrum.

#### 3.1.4. Drug release kinetics

The drug release kinetics of synthesized AgNPs was kept in a dialysis bag and released into the release medium through dialysis membrane (Fig. 3). Continuous release of silver has been observed over the period of the study (10 h). The rate of dissolution and the concentration of silver was obeyed the zero order kinetics with r^2^>0.96. The AgNPs released in to the medium were expressed as cumulative drug release % versus time (h). Throughout, the duration (10 h) of the experiment about 69% of the AgNPs were released consequently. This drug release was linked to the particle size and increasing surface area of AgNPs. Small nanoparticles underwent more rapid release kinetics than the bigger nanoparticles due to higher diffusability of the small particles. There was a difference in the final cumulative drug released throughout the study.

**Fig 3.**
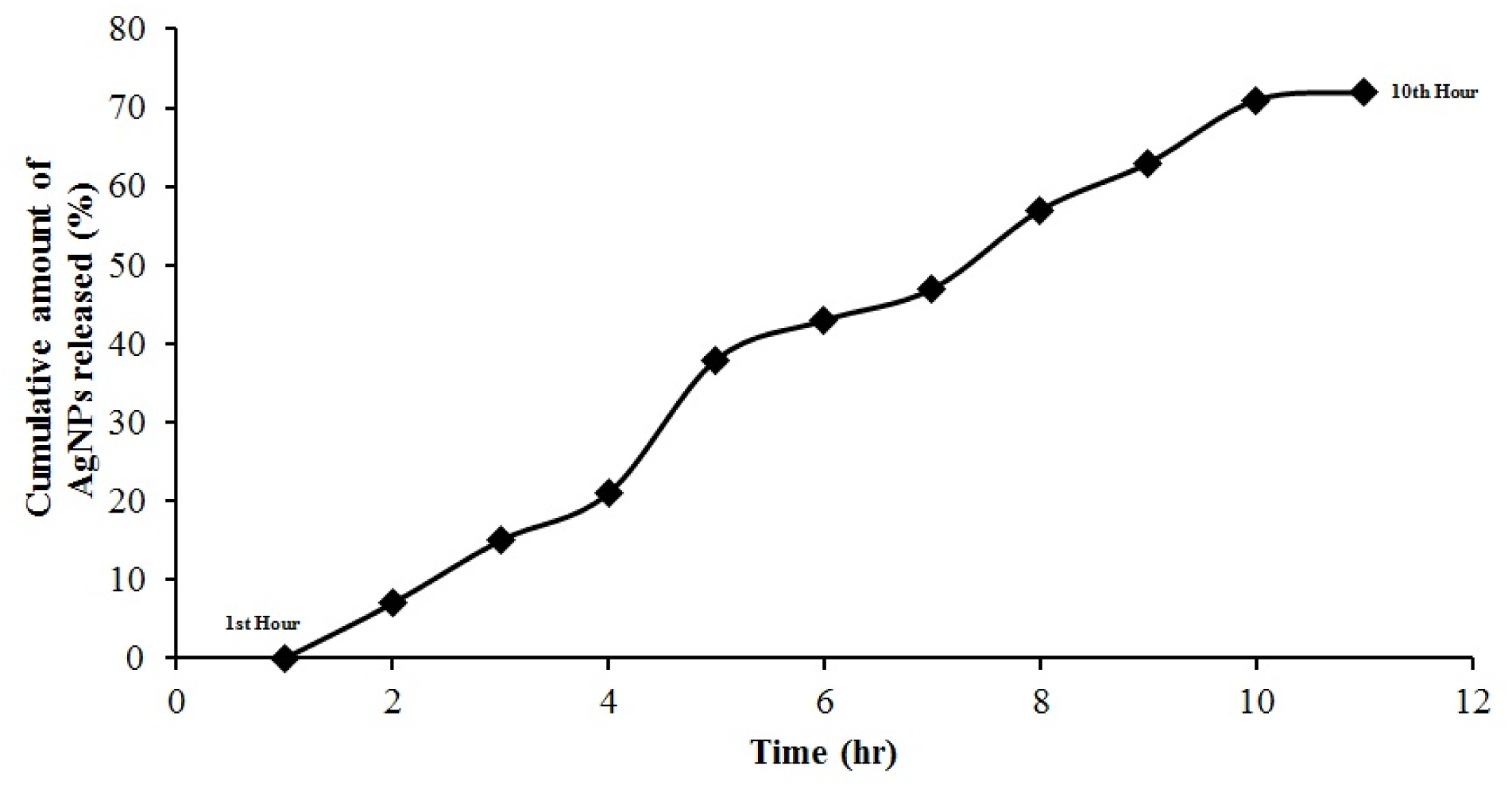
Drug release kinetics for the synthesized Ag NPs functionalized with amoxicillin drug during 10 hour duration.

**Fig 4.**
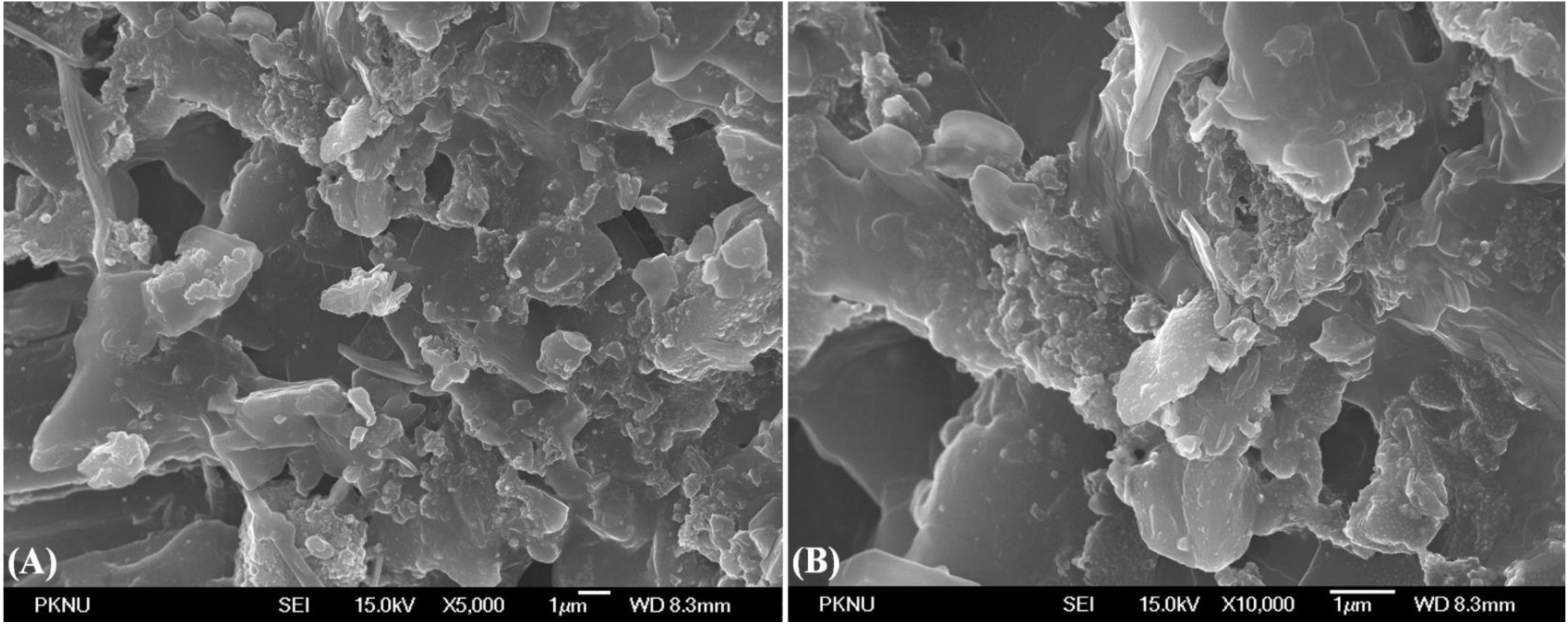
FESEM of the Ag NPs functionalized amoxicillin (A). 30,000X (B).50, 000X

#### 3.1.5. AFM and NC-AFM studies

AFM was performed to identify the topological appearance, showed with an average size of 35.50 nm, provided a zoom-in view of the surface on a nanoscale with regards to surface modification of AgNPs. The grouping of the drug and AgNPs were confirmed by the Scanning tunneling microscopy (Fig.S2 A & B). AFM analysis clearly depicted the formation of the AgNPs conjugated amoxicillin, and also that the surface morphology of the particles was uneven due to the presence of some of the aggregates and individual particles of AgNPs. Scanning probe microscopy by NC-AFM images of the AgNPs showed individual particle with an average size of 35.50nm (Fig.S3 A & B).

#### 3.1.6. Field emission scanning electron microscopy (FESEM)

Figs.4A and B depicts the FESEM image of AgNPs at 30,000 X and 50,000 X showed synthesized AgNPs were not in direct contact even within the aggregates, and indicated stabilization of the nanoparticles. The nanoparticles were distributed on the surface with the formation of aggregated and un-uniformLy nanoparticles. Particles were nano-size with smooth surface and spherical, oval and clusters. The results indicated that these surface planes having different packing structures are energetically similar to each other. Dispersed uniform spherical silver particles were prepared in the absence of protective colloid by rapidly mixing concentrated iso-ascorbic acid and silver-polyamine complex solutions. By varying the nature of the amine, temperature, concentration of reactants, silver/amine molar ratio, and the nature of the silver salt, it was possible to tailor the size of the resulting metallic particles in a wide range (80 nm to 1.3 µm) [31].

#### 3.1.7. Agar well-diffusion method

The zones of inhibition of well diffusion analysis on *E. coli* are shown in Fig.S4 A & B. The combined effect of AgNPs with amoxicillin found in the synthesized AgNPs exhibited dramatically increased the zones of inhibition. The synthesized AgNPs using amoxicillin showed significant results against *E*. *coli* cells and reported fewer efficacies when applied separately. This can be attributed to when silver nanoparticles enter the bacterial cell it forms a low molecular weight region in the center of the bacteria to which the bacteria conglomerates thus, protecting DNA from the silver ions. The nanoparticles preferably attack the respiratory chain, cell division finally leading to cell death. The nanoparticles release silver ions in the bacterial cells, which enhance their bactericidal activity [32, 33]. [34] proposed that nanoscale size and the presence of a (111) plane combine to promote the biocidal property, which belongs to silver.

#### 3.1.8. MTT Assay

The viability of cells was assessed by MTT assay using Hep-G2 cell lines [22] and the results indicated that the IC_50_ value of 50.78µg/mL. The viability of the cells was found to be increasing proportional to the concentrations of synthesized AgNPs in Fig.5. The cell populations and the capacity of proliferation decreased significantly in the samples treated with concentrations of the sample ranging from 25 to 500µg/mL in Fig.S5. Silver nanoparticles fabricated in HEPES buffer exhibited potent cytoprotective and post-infected anti-HIV-1 activities toward Hut/CCR5 cells [35]. [36] showed that the mixed-ligand silver (I) complex of 4-oxy-3-nitro-coumarin-bis (phenanthroline), could decrease the proliferation of all four cell lines including neoplastic renal and hepatic, namely A-498 and HepG2 cells, respectively, along with two non-neoplastic renal and hepatic cell lines, HK-2 and Chang, respectively.

**Fig 5.**
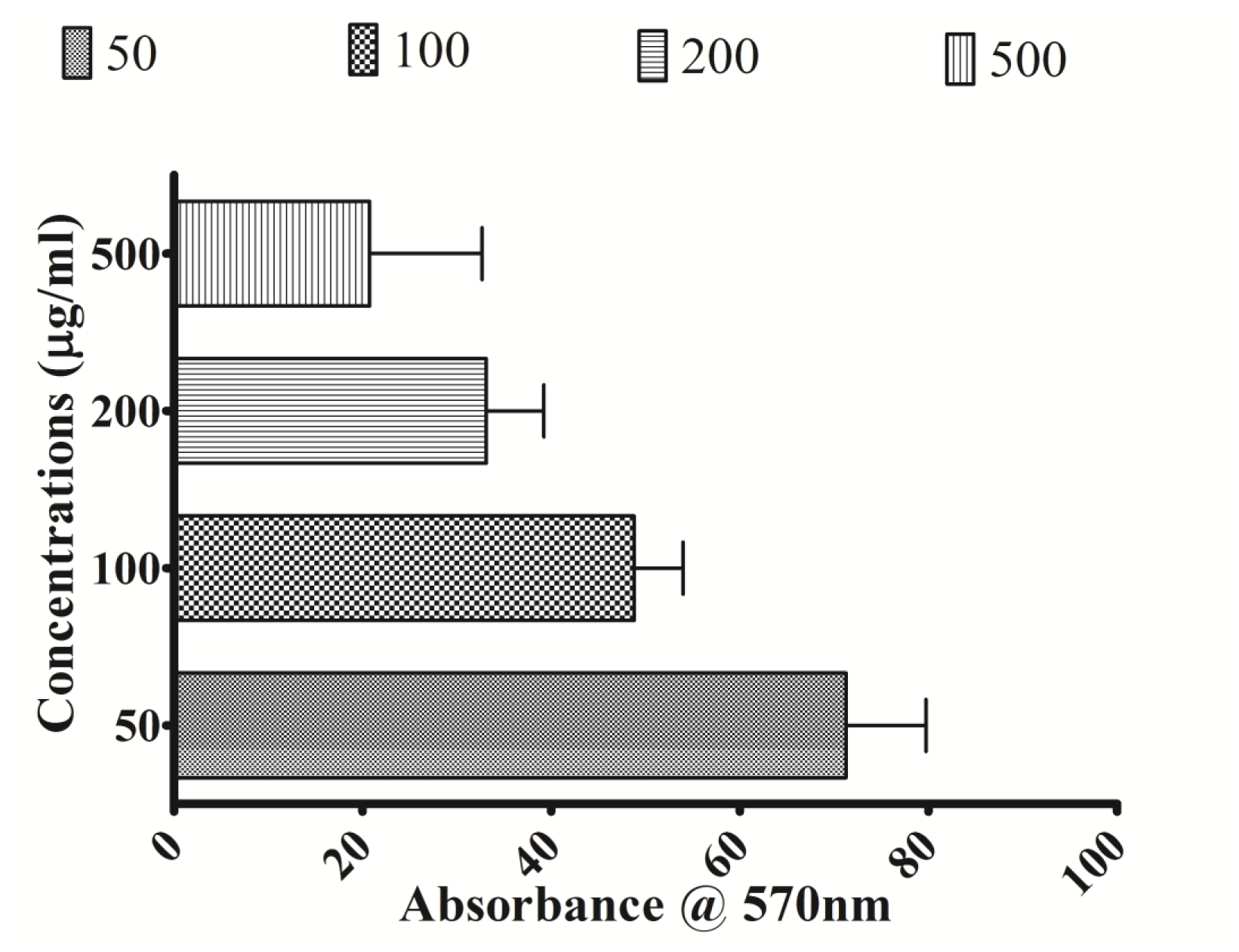
Percentage inhibition of Hep-G2 cancer cell lines against Synthesized Ag NPs.

## 4. Antioxidant activities

### 4.1. Total antioxidant activity

Total antioxidant activity of functionalized AgNPs was expressed as the number of equivalents of ascorbic acid. The phosphomolybdenum method was based on the reduction from Mo (VI) to Mo (V) by the antioxidant compound and the formation of green phosphate/ Mo (V) complex with maximum absorption observed at 695 nm [37]. The present study revealed that the antioxidant activity of AgNPs was in the increasing trend with increasing the concentration of ascorbic acid (control) and AgNPs. Among the tested AgNPs possessed potential antioxidant activity as compared with ascorbic acid (Fig. S6A).

### 4.2. Reducing power activity

The reducing power of synthesized AgNPs containing amoxicillin was a reducing agent (Fig. S6B). The presence of antioxidants caused the reduction of the Fe3^+^/ferricyanide complex to the ferrous (II) form, which could be measured by the formation of Perl’s Prussian blue at 700 nm. Ferric ion reduction was often used as an indicator for electro-donating activity of antioxidant, and reported an important mechanism of phenolic antioxidant action [38]. It was proved that when no hydroxide ion was added to the system, the Ag^+^ would not be reduced by dextrose at all, hence indicating the necessity of OH^−^ to this reduction reaction, confirming the requirement for free hydroxyl groups [39]. The reducing activity of the test sample increased steadily with their concentrations increasing.

### 4.3. DPPH radical scavenging activity

DPPH is stable nitrogen centered radical and has been widely used to test the free radical scavenging ability of various samples. The reduction capability of DPPH was determined by the decrease in its absorbance at 517 nm, which is induced by antioxidants. Positive DPPH test suggests that the samples were free radical scavengers. The DPPH radical scavenging effect of AgNPs is shown in Fig. S6C. The percentage DPPH scavenging activity ranged from 20.2% to 50.4%. The higher activity of weak antioxidants in contrast to relative ineffectiveness of potent antioxidants, in preventing Ag NP induced toxicity indicated the free radicals was not played a role in cytotoxicity. The cytotoxicity of AgNPs was mediated by free radical independent mechanisms [40]. It has been demonstrated that Ag NPs were slowly released Ag ions over duration of few days. These Ag ions were responsible for antimicrobial activity of Ag NPs and may be involved in cytotoxicity [41, 42]. Other studies have suggested that AgNPs were cytotoxic because of the oxidative stress that is independent of the toxicity of silver ions [43].

The AgNPs was noted to induce elevated levels of oxidative stress, glutathione depletion and damage to the cell membrane as found from the adenylate kinase assay and that leads to the apoptosis. Overall, significant differences were observed between the sensitivity of the two cell lines which could be understood in terms of their natural antioxidant levels [44]. This consequently may enhance their absorption when applied *in vivo*. The above results suggest that the AgNPs would be stable and rapidly released when applied at the infection sites. The present results suggested that synthesized AgNPs with amoxicillin showed significant antibacterial activity. The significant points suggesting the increase in the antagonistic activity of the synthesized AgNPs were to be ascribed by the rate of dissolution and the concentration of silver obeyed the zero order kinetics with r^2^>0.96. The drug release was linked to the particle size and increasing surface area of the AgNPs. The grouping of the drug and AgNPs were confirmed by the Scanning tunneling microscopy. The viability of the cells was found to be increasing proportional to the concentrations of the synthesized Ag. The cell populations and the capacity of proliferation decreased significantly in the samples treated with concentrations of the sample ranging from 25 to 500µg/mL. This article applies elements of the drug delivery paradigm to nanosilver from amoxicillin as a dissolution agent and presents a methodical study of chemical notions for controlled release.

## 5. Conclusion

To conclude, we have used non-toxic, eco-friendly way for avoidance of the drug resistance by the combined treatment of AgNPs with amoxicillin. The presence of organic compounds with a carboxyl moiety was confirmed on the surface of the AgNPs fabricated by this combined organic and inorganic substances, which we have designated as a semi-biosynthesis method. Metal nanoparticles combined with antibiotics may therefore show improved efficacy in specific treatment, thus lowering side effects associated with drug molecules. The surface chemistry of the as prepared AgNPs functionalized was studied using infrared spectroscopy.

## 6. Acknowledgement

The authors are grateful to acknowledge the support extended by SAIF, Cochin University of Science and Technology, Cochin for characterization of nanoparticles.

