## Supplementary File for "‘Synergistic-cidal’ effect of amoxicillin conjugated silver nanoparticles against *Escherichia coli*"

### Antibacterial, antioxidant and cytotoxic activities of amoxicillin conjugated silver nanoparticles

**Supplementary information**

**Fig.S1**.Hypothetical plan for the formation of synthesized Ag NPs nanoparticles functionalized amoxicillin

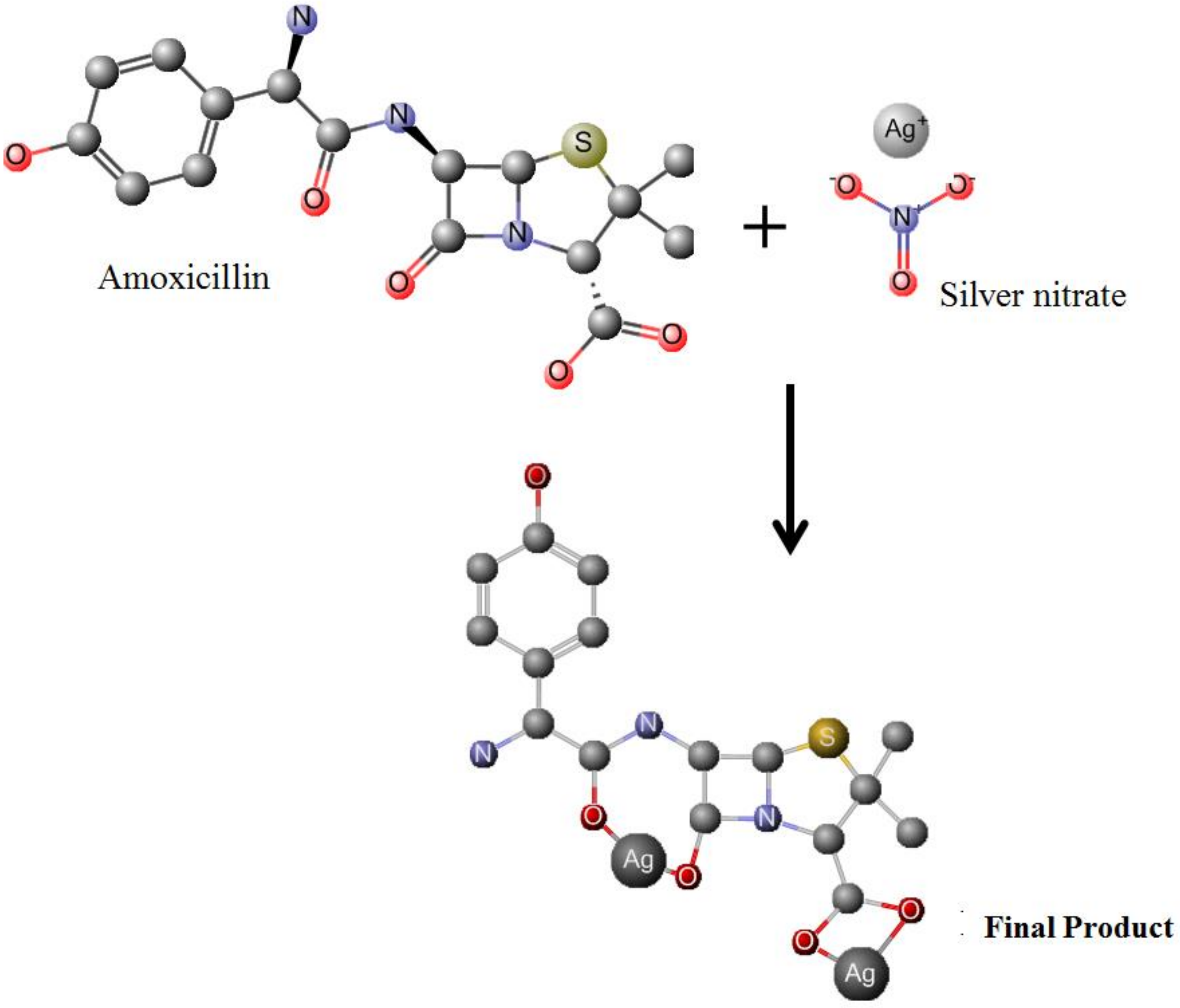

**Fig.S2.** AFM image of the silver nanoparticles functionalized amoxicillin showing increased surface area (A) Synthesized Ag NPs (B) Single particle view.

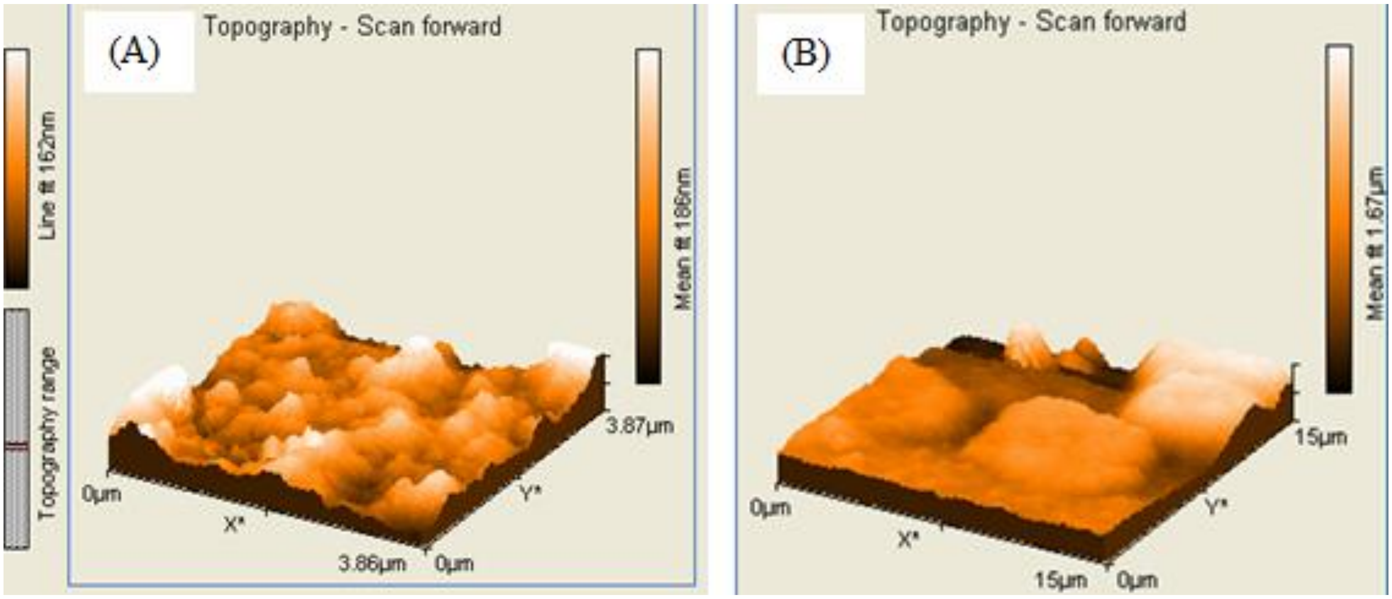

**Fig.S3** NC-AFM mode of the Ag NPs functionalized amoxicillin showing individual (A) Synthesized Ag NPs (B) Particle size analysis showing average size at 35.50nm.

.

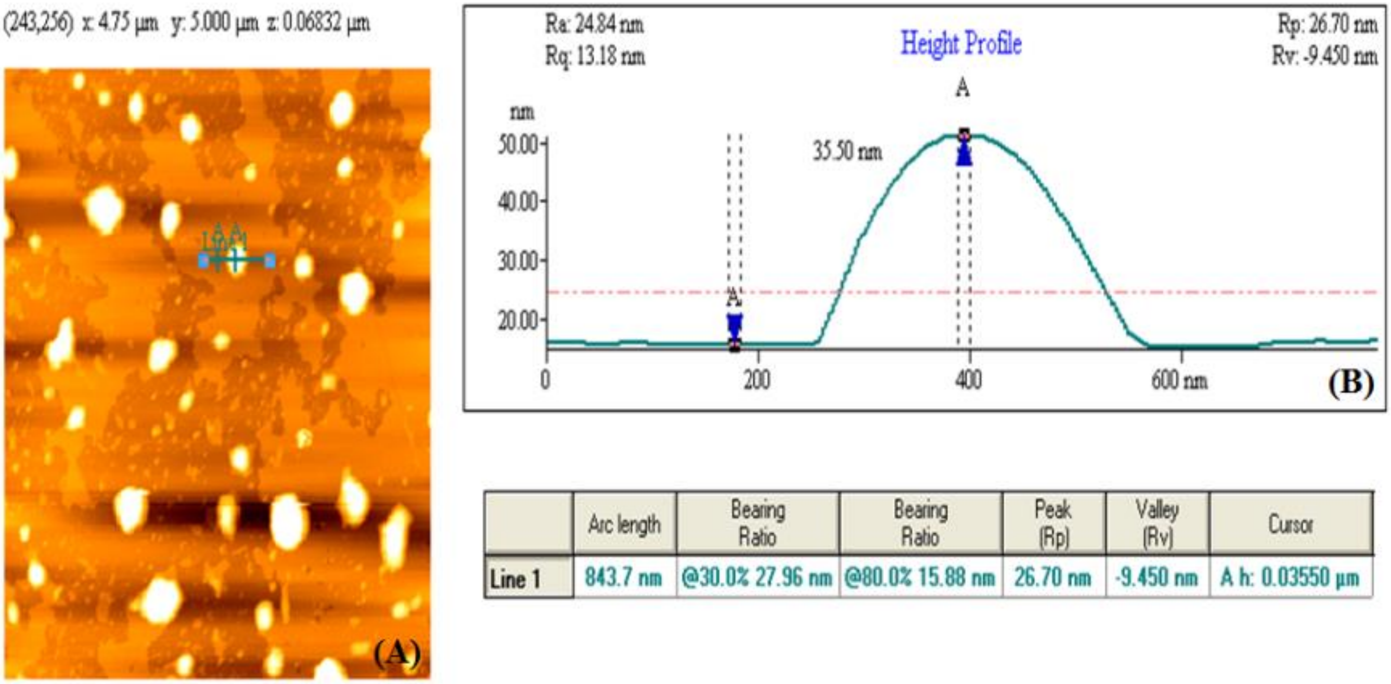


| | Arc length | Bearing Ratio | Bearing Ratio | Peak (Rp) | Valley (Rv) | Cursor |
|---|---|---|---|---|---|---|
| Line 1 | 843.7 nm | @30.0% 27.96 nm | @80.0% 15.88 nm | 26.70 nm | -9.450 nm | A h: 0.03550 µm |

**Fig.S4.** Well diffusion plates for the (A). Synthesized Ag NPs (B). Amoxicillin drug against *E. coli* showing increased zones of inhibition.

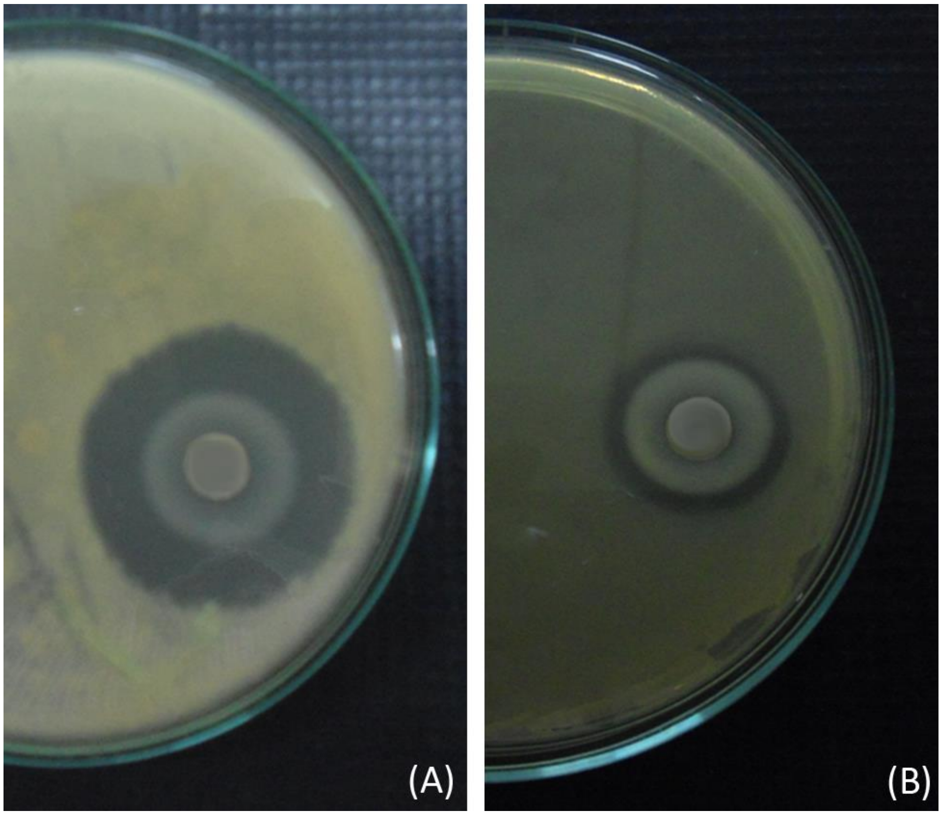

**Fig.S5**. Anti-cancer activity of Synthesized Ag NPs against Hep-G2 cancer cell lines (A). 50µg/ml (B). 100 µg/ml (C). 200µg/ml (D). 500 µg/ml

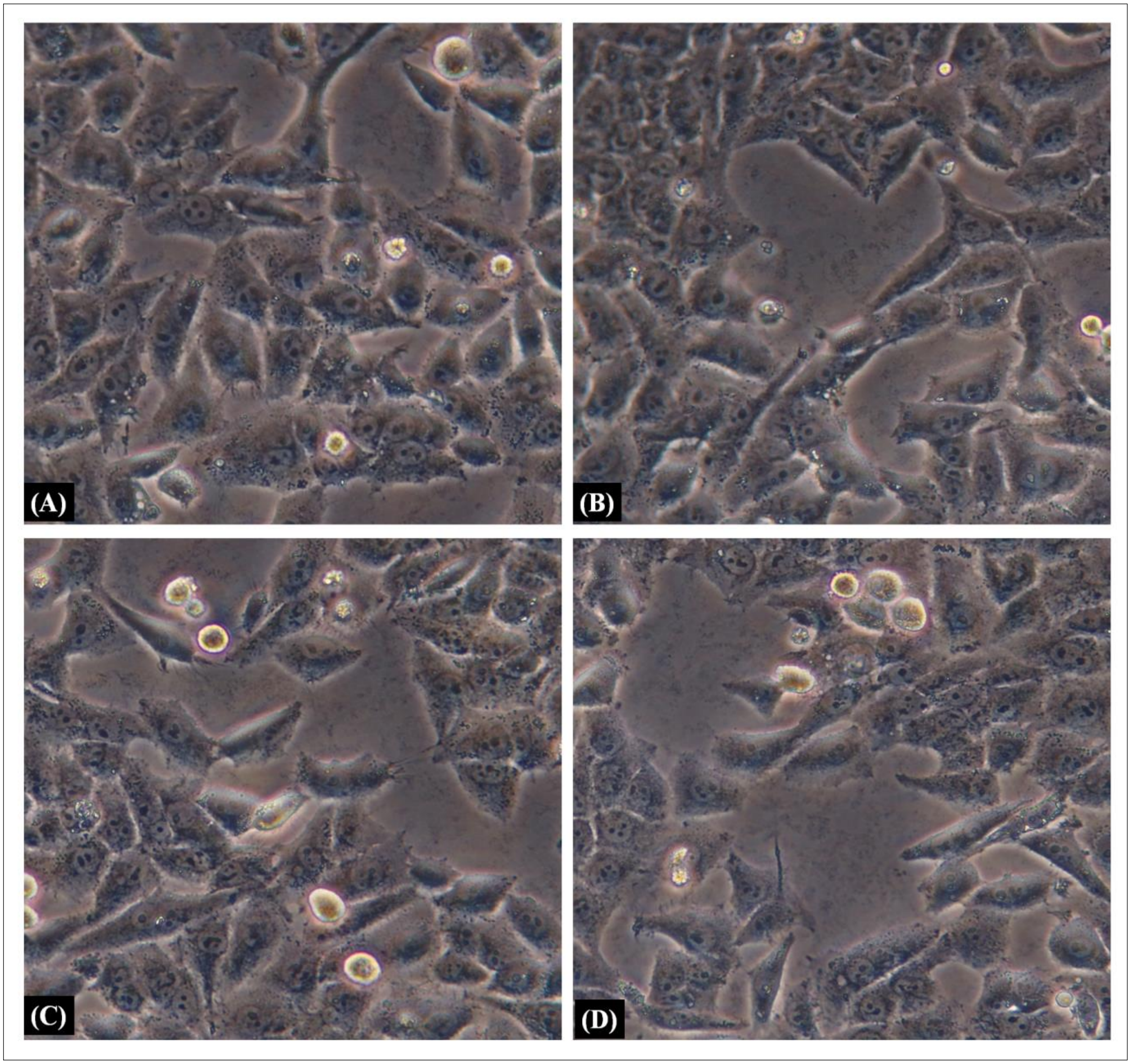

**Fig.S6** Antioxidant activity of the synthesized Ag NPs (A). Total Antioxidant Activity (B). Reducing power activity (C). DPPH radical scavenging activity

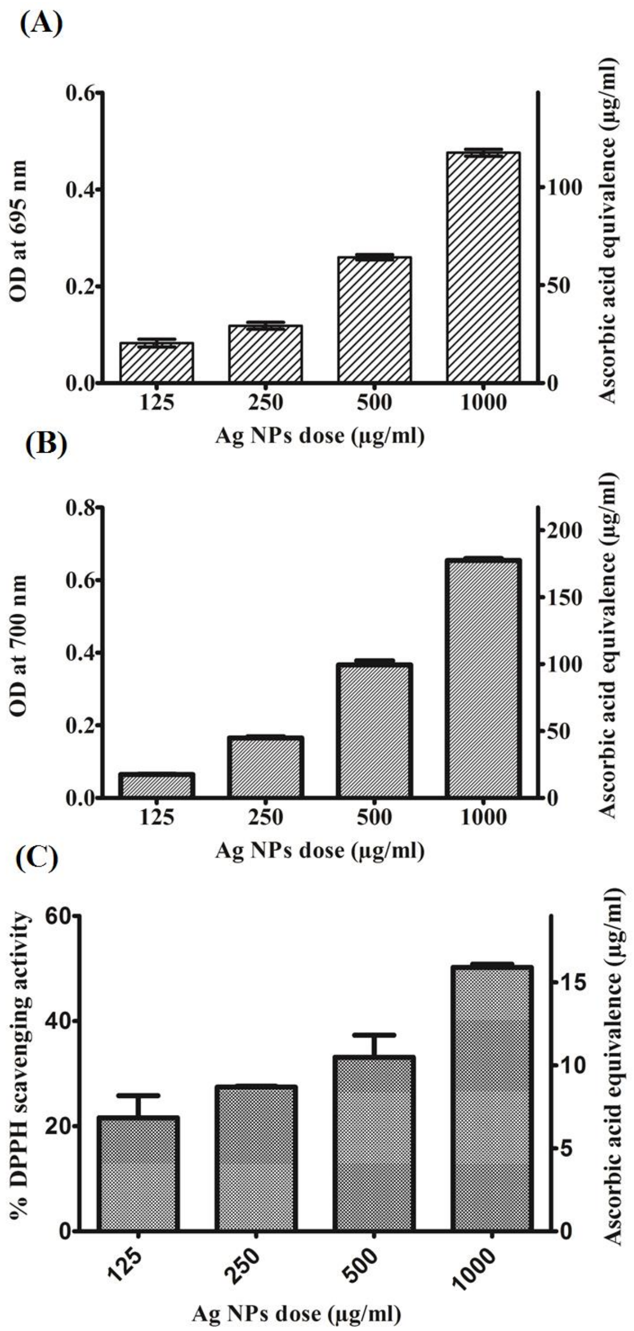
